# The Mammalian KU70 C-terminus SAP Domain Is Required to Repair Exogenous DNA Damage

**DOI:** 10.1101/2024.06.30.601420

**Authors:** Yuan Wang, Michael S. Czap, Hailey Kim, Huimei Lu, Jingmei Liu, Yokechen Chang, Peter J. Romanienko, Cristina Montagna, Zhiyuan Shen

## Abstract

The mammalian non-homologous end joining (NHEJ) is required for V(D)J recombination as well as coping with exogenously induced DNA double strand breaks (DSBs). Initiated by the binding of KU70/KU80 (KU) dimer to DNA ends and the subsequent recruitment of the DNA-dependent protein kinase catalytic subunit (DNA-PKcs), NHEJ plays a key role in DNA repair. While there has been significant structural understandings of how KU70 participates in NHEJ, the specific function of its highly conserved C-terminal SAP domain remains elusive. In this study, we developed a novel mouse model by deleting the SAP domain but preserving the KU70 nuclear localization and its dimerization ability with KU80. We found that the KU70 SAP deletion did not affect the V(D)J recombination or animal development but significantly impaired the animals and cells in repairing exogenously induced DSBs. We further showed an inability of KU70-ΔSAP cells to retain the DNA Ligase IV (LIG4) and other NHEJ co-factors on chromatin, and a spreading pattern of DSB marker γH2AX in KU70-ΔSAP cells after DNA damage. Our findings suggest that a specific inhibition of the SAP function may offer an opportunity to modulate cell sensitivity to therapeutic DSB-inducing agents without interfering with the developmental function of KU70.

**KeyPoints:**

- Generation of a novel transgenic mouse line lacking the C-terminal conserved KU70-SAP domain
- KU70-SAP defends against exogenous DSBs, but unessential for development and V(D)J recombination
- KU70-SAP aids in recruiting and retaining NHEJ components, such as LIG4, to DSB sites

## Introduction

Double strand breaks (DSBs) are the most lethal form of DNA damage that can be induced by ionizing radiation (IR) and radiomimetic agents. Efficient and precise DSB repair is critical for cell survival and the stability of their genomes. DSBs are mainly repaired by the non-homologous end joining (NHEJ) and homologous recombinational repair (HRR) machineries. In mammalian cells, NHEJ defects have more profound sensitivity to IR than HRR defects (1,2), reflecting the prominent role of NHEJ in repairing exogenously induced DSBs. It has long been known that the end-loading and encircling of the double stranded DNA (dsDNA) through the aperture-like structure formed between the KU dimer are the initiating events for NHEJ (3).

However, it is until recent years when several structural analyses have collectively painted an elegant picture on how a KU-initiated large protein complex evolves to align two DSB ends for rejoining (4–8). The DNA-bound KU dimers recruit DNA-PKcs and then slide inward for about 15 nt to allow DNA-PKcs to occupy the DNA ends (9). Recruitments of additional proteins including LIG4, X-ray repair cross-complementing 4 (XRCC4) and XRCC4-like factor (XLF) form a long-range synapsis protein complex to hold two DNA ends along with two DNA-PK complexes in proximity (10). After the synapsed DNA ends are processed and become ligation-compatible, autophosphorylation of DNA-PKcs results in its eviction to transform the long-range synapsis complex to a short-range one that brings the two DSB ends to a close proximity for ligation by an iterative mechanism (6). Later, a presumable LIG4 re-configuration allows its ligase domain to reach the dsDNA ends and ligate, one strand at a time by each of the two LIG4 molecules in the short-range synapsis complex (4–6).

In human, the KU70 subunit is a protein with 609 amino acids (aa). The N-terminal 538 aa is responsible for forming a stable heterodimer with KU80 and the C-terminal 71 aa consist of a flexible linker region (aa 539-558) that contains the nuclear localization signal (NLS), and a globular domain of 51 aa (aa 559-609) that was historically referred as the KU70-SAP (SAF- A/B, Acinus, and PIAS) domain (11). It is noteworthy that only the C-terminal 37 aa (referred as cSAP, aa 573-609) of the KU70-SAP was originally defined as the canonical SAP motif (12), but the 14 aa helical domain (referred as nSAP, aa 559-572), unique for KU70, may also contain a weaker NLS activity (Figure S1).

In some of the crystallography and CryoEM studies (6,10), the C-terminal linker and KU70-SAP were found to be absent in the DNA-PK complex, likely due to the flexibility of the linker region and mobility of the KU70-SAP domain. Others were able to capture at least three different KU70-SAP positions in the DNA-bound and DNA-free forms of the KU dimer (7,8,13). Initially, the KU70-SAP was shown to associate with the α/β-domain of KU80 of the DNA-free KU dimer (7,8). Using a full length KU70 and C-terminally truncated KU80 (trKU80), it was recently shown that the KU70-SAP domain can also locate near the aperture of the DNA-free KU dimer, but may move away from the aperture to an area near aa 450-538 of KU70 after the KU70/trKU80 dimer binds to DNA (13). These works suggested that the KU70-SAP may change positions in a dynamic NHEJ complex depending on the stage of the repair process, indicating a regulatory or coordinating role of the KU70-SAP domain. Interestingly, in all these scenarios, it was the nSAP helical domain that was involved in interacting with the main body of the KU dimer at various positions (7,8,13).

Given its ability to bind RNA and DNA (14–16) and the conservative feature of the SAP sequence, there is substantial conjecture surrounding the essential role of SAP domain in KU70- associated processes. However, evidence regarding its specific role in KU70 function remains inconclusive, as structural analyses have yet to offer a definitive confirmation or refutation.

Engineered deletion of the plant *Arabidopsis thaliana* at KU70-SAP (aa 593-621), corresponding to human aa 581-609 of the cSAP, resulted in decreased KU loading to DNA ends *in vitro*.

However, this alteration did not have an impact on classical NHEJ, telomere maintenance, or plant development (15).

To directly investigate the biological consequence of mammalian SAP deletion, we used CRISPR technique to generate a novel transgenic mouse model in which the KU70-SAP domain (aa 564-608) was deleted while the flexible linker with NLS was retained. We found that deletion of the KU70-SAP did not impact the stability of KU80 protein, animal growth, or the V(D)J recombination demonstrated through class-switch recombination in the B cells. However, it did sensitize animals to IR-induced BM failure, elevated spontaneous and IR-induced chromosome breakages and aberrations, heightened cells sensitivity to DSB-inducing chemotherapy agents, reduced KU70 recruitment to DNA damage sites, diminished LIG4 retention and recruitment of other NHEJ factors to chromatin. Our findings suggest a critical role of the SAP domain in repairing exogenous DNA damage, particularly distinct from the physiological substrates like DSBs encountered during immunoglobulin class-switch. Our study represents a pioneering investigation confirming the significant biological role of the KU70-SAP domain in the repair of exogenous DNA damage, likely mediated through a dynamic regulation of the LIG4 activity at the DSB sites. Moreover, it further indicates that targeting the SAP-related function(s) of KU70 could be a viable strategy to sensitize mammalian cells to therapeutic DNA damage with minimum impact on the endogenous and physiological function of KU70.

## Materials and Methods

### Mouse line and total body irradiation (TBI)

A KU70-A564* knock-in (KI) mouse line (ΔSAP) was generated using a single gRNA and a mixture of two templates adjacent to the Cas9 cutting site. Briefly, the wild-type (WT) mouse KU70 (mKU70) nucleotide and amino acid sequences starting from A564 were modified by changing GCCC to tGAAC. This resulted in the insertion of one nucleotide and the creation of a stop codon, terminating mKU70 at A564. Additionally, this introduced a new XmnI (GAANN^NNTTC) restriction site, which was highlighted in blue in Figure 1A and used for genotyping. Mice were housed in individually ventilated cages within a specific pathogen-free facility, maintained on a 12-hour light/dark cycle, and provided with ad libitum access to food and water. All animal work presented in this study was approved by the Institutional Animal Care and Use Committee (IACUC) at Rutgers Robert Johnson Medical School. We adhered to institutional guidelines regarding animal welfare. For TBI experiments, mice were temporarily restrained in the ventilated pie-shaped chambers of the RadDisk cage. This cage is circular with 8 pie-shaped chambers and a rotating lid that covers 7 of the chambers. Up to 7 mice were individually placed in the separate chambers of the RadDisk cage every time, which was then placed inside the Gammacell 40 Extractor (MDS Nordion) 137Cs γ-irradiator. Specific radiation doses are provided in the relevant experimental descriptions.

**Figure 1.**
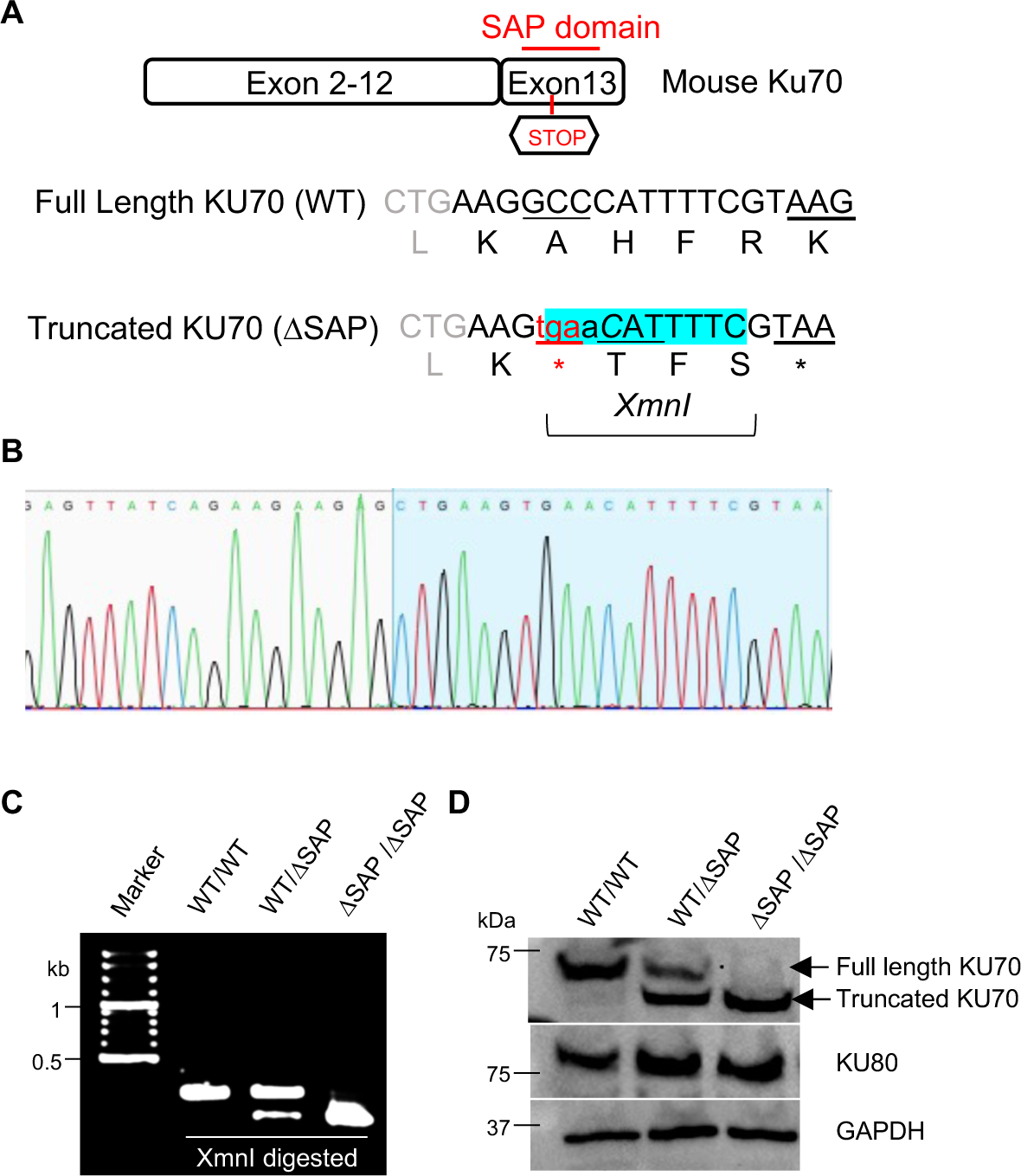
**Generation and verification of KU70-**Δ**SAP mouse line.** (**A**) Scheme of the CRISPR-Cas9 editing strategy for mouse Ku70 gene. The upper sequence indicates the WT mouse KU70 nucleotide and amino acid sequences starting from L562, and the lower sequence indicates A564* mouse line (ΔSAP) in which GCCC was changed to tGAAC, resulting in one nucleotide insertion and the creation of a new STOP codon to terminate the mouse KU70 at A564. This also generates a new XmnI (GAANN^NNTTC) site highlighted in blue to be used for genotyping. (**B**) ABI sequencing chromatogram confirming the insert of a STOP codon. DNA from homozygous KU70-ΔSAP mouse was used for ABI Sanger sequencing. (**C**) Agarose gel electrophoresis of PCR-amplified products with XmnI digestion from WT and ΔSAP mice. (**D**) The expression of full length and truncated KU70 in MEFs which from indicated KU70 KI mouse lines, KU80 was detected as the binding partner of KU70. GAPDH was used as the loading control.

### PCR genotyping and XmnI digestion

The genomic DNA (gDNA) was extracted from 1-2 mm tail tissue and incubated with a lysis buffer (25 mM NaOH, 0.2 mM EDTA) in a PCR machine for 30 minutes at 95°C, followed by cooling to room temperature (RT). An equal volume of neutralization solution (40 mM Tris-HCl, pH 5.0) was then added, and the sample was mixed by shaking several times. For the PCR reactions, 1 µL of the gDNA was used in 10 µL reactions with mKU70 PCR primers: 5’- TGTAGCTGGGCTATCACTGGAAGC-3’ (forward) and 5’-CTTTGTAGCTACCACCTAGCTTGG-3’ (reverse), which flank a region containing a newly introduced XmnI digestion site. PCR amplification was performed using a PCR premix solution (Syd Labs, MB067-EQ2B-L). Following PCR, the product was digested with the XmnI enzyme (NEB, R0194L) at 37°C for 3 hours. The digested PCR product was then prepared for agarose gel electrophoresis.

### Collection of Lineage negative (Lin^-^) cells and flow cytometric analysis

Bone marrow (BM) single-cell suspensions were prepared from either one non-irradiated mouse or two irradiated mice, including the tibia and femur pair and the spine for hematopoietic stem cell (HSC) subset analysis. Following bone crush in RPMI1640 medium supplemented with 10% FBS, 1% Glutamine, 1% Penicillin/Streptomycin, and 0.01% 2-Mercaptoethanol, red blood cells (RBCs) were lysed using RBC lysis buffer (Sigma-Aldrich, R7757). For lineage depletion, the cells underwent lineage cell depletion using the lineage cell depletion kit (Miltenyi Biotec, #130- 090-858) following the manufacturer’s instruction. Briefly, the cell pellet was resuspended in FACS buffer (PBS with 1% BSA) containing 20 μL of Biotin-antibody cocktail for 10 minutes, followed by the addition of 40 μL of anti-biotin microbeads for another 15 minutes on ice. After incubation, cells were washed once with FACS buffer and then loaded onto a pre-wet LS column (Miltenyi Biotec, #130-042-401) on the MACS Multistand (Miltenyi Biotec, #130-108-936) using the magnetic MidiMACS separator (Miltenyi Biotec, #130-042-302). The Lin^-^ cells were collected from the effluent. For flow cytometric analysis, cells were stained using Fixable Yellow Live/Dead Dye (eBioscience, #65-0866-14) and specific antibodies as described (17).

Following staining, cells were washed twice with FACS buffer and prepared for flow cytometry (Cytek Aurora Analyzer, SKU IMSR-4.6a). FlowJo^TM^ portal (v10.9.0) was used for the analysis. UltraComp ebeadsTM compensation beads (Invitrogen, #01-2222-42) were used for compensation setup.

### Mouse splenic B cell isolation and activation

The mouse spleen was placed into a 60 mm petri dish containing B cell wash buffer (1x Hanks with 1% heat-inactivated FBS and 1% penicillin-streptomycin) and minced into small pieces using a razor blade. The tissue along with the buffer was then transferred into a 70 μm strainer within a 50 mL tube. The spleen fragments were further gently disrupted within the mesh using the flat end of a syringe, and the cells were collected from the 50 mL tube following centrifugation at 1,200 rpm for 10 minutes. The resultant cell pellet was treated with ACK lysing buffer (Gibco, #A1049201) and neutralized by adding wash buffer, followed by another round of centrifugation. The cell pellet was gently tapped to loosen it and then resuspended in 1 mL of wash buffer containing 50 μL of CD43 (Ly-48) MACS microbeads (Miltenyi Biotec, #130-049- 801). The mixture was incubated on ice for 30 minutes. After incubation, the cells were washed by adding wash buffer, gently mixing, and centrifuging at 1,200 rpm for 10 minutes. The sample was resuspended in 1 mL of wash buffer and B cells were separated from the effluent fraction using a MidiMACS™ Separator with an LS column, as described in the Lin^-^ cell collection section. The column was washed twice, and the appropriate cells were placed in a 6-well plate or flask containing B cell culture media (RPMI 1640, 10% heat-inactivated FBS, 1% penicillin-streptomycin, 1% glutamine, 1x nonessential amino acids, 1% sodium pyruvate, 53 mM 2- mercaptoethanol, and 10 mM HEPES) supplemented with 25 μg/mL LPS (Sigma, L2630) and 5 ng/mL IL-4 (Sigma, I1020). The cells were cultured for 72 hours at 37°C in a 5% CO_2_ incubator.

### Class switch recombination assay

The B cells were collected at 72 hours post-isolation and activation, and then washed once with cold PBS. The cells were blocked with anti-CD16/32 antibody (BD Biosciences, #553141) at RT for 10 minutes and stained with FITC anti-B220 (BD Biosciences, #553088), PE anti-IgM (BD Biosciences, #562033), and Biotin anti-IgG1 (BD Biosciences, #553441) antibodies at 4°C for 30 minutes. Following staining, the cells were centrifuged and washed twice with PBS, then stained with Streptavidin Alexa Fluor^TM^ 647 conjugate antibody (Invitrogen, S32357) for an additional 10 minutes at RT. Flow cytometry was then used for analysis.

### Chromosome metaphase spread analysis

The B cells were irradiated at 2 Gy, 72 hours post-isolation and activation, using 7.5 μg/mL Concanavalin A (Sigma, C5275). Following a 12-16 h incubation post-IR, the cells were cultured in the presence of 10 μg/mL Colcemid solution (Roche, #295892) for 30 minutes to enrich for metaphase cells. The metaphase cells were collected, swollen in pre-warmed 0.075 M KCl at 37°C for 25 minutes, and then fixed in a freshly prepared 3:1 mixture of methanol and acetic acid. Cell preparations were dropped onto slides and allowed to dry overnight. The slides were then stained using a DAPI Fluoromount G medium (SouthernBiotech, #0100-20) for visualization. Metaphase images were acquired with a BioView scanning system mounted on and Olympus BX63 microscope (BioView USA Inc., Billerica, MA) with fine focusing oil immersion lens (×60, NA 1.35 oil). Multiple focal planes (n = 11, 0.5 µm per plane) were acquired to ensure that scanning on different focal planes was included.

### Generation of mouse embryonic fibroblasts (MEFs) from KU70-ΔSAP mouse model

KU70-ΔSAP heterozygous mice were bred at 3 months of age, and upon confirmation of pregnancy, female was euthanized to harvest E15.5 embryos. The MEFs were generated as previously described (18). Briefly, embryos were extracted from the uterus. The skin from the back of each embryo was removed, minced, and subjected to enzymatic digestion in 0.25% Trypsin-EDTA solution at 37°C for 30 minutes. Following digestion, α-Minimum Essential Medium (α-MEM) with 10% fetal bovine serum (FBS) was added to inactivate the trypsin. The mixture was gently agitated, and the cell suspension was left undisturbed for about 5 minutes to allow larger embryo fragments to settle. The supernatant containing single cells was then transferred to a new tube and centrifuged at 1,000 rpm for 3 minutes. The resulting cell pellet was resuspended in α-MEM supplemented with 10% FBS, 20 mM glutamine, and 1% penicillin-streptomycin, and the cells were cultured at 37°C in a 5% CO_2_ incubator.

### Micro-irradiation and Immunofluorescence (IF) staining

MEFs were seeded in chamber slides with 1 μM IdU one day prior to micro-irradiation. Laser micro-irradiation was performed using a PALM MicroBeam Laser-capture Microdissection System (Germany) equipped with a Zeiss Axio observer microscope (Carl Zeiss AG) on a custom-designed granite plate. A LabTek chamber slide with live cells was mounted on the microscope stage integrated with the PALM Microlaser workstation. The cells were visualized under visible light, laser-targeted nuclei were selected using a Zeiss software (PALM Robo V4.9), and the nuclei were subsequently irradiated with a pulsed solid-state UV-A laser coupled to the bright-field path of the microscope focused through an LD 40x objective lens. The laser settings were optimized to generate DNA damage restricted to the laser path, dependent on pre-sensitization, with minimal cellular toxicity. At the indicated time points post-micro-irradiation, cells were fixed with Formalde-Fresh Solution (Fisher, SF93-4) for 15 minutes and then permeabilized with 0.15% Triton X-100 for 15 minutes. After three washes with PBS, the fixed cells were incubated with primary antibodies in 2% BSA blocking buffer at RT for 1 hour, followed by incubation with FITC or TRITC conjugated anti-mouse or anti-rabbit secondary antibodies at RT for 1 hour. Nuclei were visualized by DNA staining with DAPI Fluoromount G medium. For KU70 staining, cells were incubated for 5 minutes at RT with CSK buffer (10 mM Pipes, pH 7.0; 100 mM NaCl, 300 mM sucrose, and 3 mM MgCl_2_) containing 0.1% Triton X- 100 and 0.3 mg/mL RNase A before the fixation step.

### Plasmids and production of lentiviruses and retrovirus

To achieve endogenous KU70 inducible knockdown, KU70 shRNA targeting the 3’-UTR of the hKU70 gene was cloned into the pLKO-Tet-Neo vector, which confers neomycin resistance. For lentivirus production, 293T cells were co-transfected with pLKO-shKU70-Neo, psPAX2, and pMD2G at a 2:1:1 ratio. Seventy-two hours post-transfection, the virus-containing supernatant was collected and filtered through a 0.45 μm nylon mesh and used to infect H1299 cells in the presence of 8 μg/mL polybrene. The infection process was repeated twice. Following infection, the cells were allowed to recover overnight in fresh medium and were then subjected to selection with 800 μg/mL neomycin for 7 days. The WT and truncated cDNA of hKU70 were cloned into the pLXSP-Myc-EGFP vector, which provides puromycin resistance. The DNA constructs were transfected into Phenix A cells to generate retrovirus. H1299-shKU70 cells were then infected with the produced retrovirus through three infection cycles (8 hours of infection followed by 16 hours of incubation with fresh medium per cycle). Post-infection, the cells were subjected to selection with 1 μg/mL puromycin. Stable cell lines were established by isolating and expanding positive single clones. The cells were cultured in α-MEM supplemented with 10% FBS, 20 mM glutamine, and 1% penicillin-streptomycin at 37°C containing 5% CO_2_ incubator.

### Antibodies

The following primary antibodies were used: KU70 (Invitrogen, MA5-42413; Santa Cruz, sc- 56129), KU80 (Santa Cruz, sc-5280), GAPDH (Santa Cruz, sc-365062), γH2AX (Santa Cruz, sc- 517348; Cell Signaling Technology, #2577L), Myc (Santa Cruz, sc-40); XRCC4 (Santa Cruz, sc-365118), XLF (Santa Cruz, sc-393844), LIG4 (Santa Cruz, sc-28232; Cell Signaling Technology, #14649S), Total DNA-PKcs (Cell Signaling Technology, #12311S), phosphorylated DNA-PKcs (S2056) (Abcam, ab-18192-100), Histone H4 (Cell Signaling Technology, #2935T).

### Chromatin fractionation

H1299-shKU70 cell lines stably expressing Myc-EGFP-tagged KU70 (WT and mutants) were treated with 0.2 μg/mL Dox for 5 days to knock down endogenous KU70, and then treated with or without IR at 6.5 Gy. At the 30 mins post-IR, the cells were collected and incubated on ice with permeabilization buffer (10 mM HEPES, pH 7.4; 10 mM KCl; 0.05% NP-40) supplemented with protease inhibitors. Samples were centrifuged at 12,000 rpm for 5 minutes at 4°C to separate the supernatants as the cytoplasmic fractions (S1). The pellets were washed once with permeabilization buffer, then centrifuged again at 12,000 rpm for 5 minutes at 4°C. The supernatants were discarded, and the pellets were treated with nuclease reaction buffer (10 mM HEPES, pH 7.4; 10 mM KCl; 0.5 mM MgCl_2_; 2 mM CaCl_2_) supplemented with Benzonase nuclease for 30 minutes at 4°C. Samples were then centrifuged at 12,000 rpm for 5 minutes at 4°C. The resultant supernatants, representing the nuclear fractions (S2), were collected. The remaining pellets were extracted for chromatin fractions using 0.2 N HCl on ice for 10 minutes. The supernatant was neutralized with an equal volume of 1 M Tris-HCl (pH 8.0).

## Results

### Deletion of mouse KU70-SAP domain had minimal impact on animal growth, spontaneous tumor formation, and class-switch of immunoglobulins in B cells

Despite speculation about the role of the SAP domain in KU70 functions based on *in vitro* biochemical analyses (14–16), structural analysis has not provided a clear indication of whether the KU70-SAP domain is essential for the NHEJ complex, and it remains unknown whether KU70-SAP has a functional role in mammalian development and the repair of genotoxic damage. To directly address this long-standing knowledge gap, we resorted to the CRISPR knock-in approach to modify the coding sequence in exon-13 of mouse Ku70 as illustrated in Figure 1A. This resulted in a new transgenic mouse line, designated as *Ku70-*Δ*SAP*, which codes for a KU70 protein lacking the C-terminal aa 564-608 in mice, corresponding to aa 566-609 in humans. It is noteworthy to point out that this deletion included both the nSAP and cSAP regions of KU70, while preserving the linker region that contains the NLS in the truncated KU70-ΔSAP (aa 1-563 in mice, corresponding to aa 1-565 in humans). The authenticity of the homozygous founder line was confirmed by Sanger DNA sequencing of the *Ku70* locus (Figure 1B).

Additionally, the new mouse line was validated by XmnI digestion of PCR amplified DNA covering the modified region of the *Ku70-ΔSAP* line (Figure 1C). Furthermore, Western Blot analyses of proteins extracted from homozygous and heterozygous *Ku70-*Δ*SAP* mice confirmed the expression of the truncated form of mouse KU70 protein (Figure 1D). In *Ku70* null mice, abolition of the KU dimer and destabilization of KU80 were observed, along with defective immunoglobulin class switching in B cells and impairment in animal growth (19–21). However, the *KU70-*Δ*SAP* mice showed no reduction in KU80 protein levels (Figure 1D), indicating that the truncation does not destabilize the KU dimer. This result supports the idea that stabilization of the KU heterodimer relies on the dimer-forming N-terminal region of KU70, rather than the C-terminal SAP domain (7,8). In contrast to the previously reported phenotypes of smaller body size, shorter lifespan, and spontaneous T-lymphoma development in *Ku70* null mice (19,22), the *Ku70-*Δ*SAP* mice exhibited normal body weight (Figure S2A), unchanged overall survival within the observed ∼600 days of age, and insignificant change of spontaneous tumor incidence (Figure S2B).

The *Ku70* null mice exhibit V(D)J recombination defects, including issues with class-switching in IgM positive B-cells (20,23). To determine whether the KU70-SAP domain influences lymphocyte development and the conversion of IgM to IgG1, we isolated splenic B cells from the KU70-ΔSAP KI mouse line. After *in vitro* stimulation, we compared the expression levels of IgG1 from IgM between KU70 WT and KU70-ΔSAP mice. Despite the anticipated role of KU70-SAP domain in the NHEJ, our results showed that its deletion did not affect the conversion from IgM to IgG1 in mouse splenic B cells, as compared to their sex-matched WT littermates (Figure S2C), indicating a minimum role of KU70-SAP in the class switch V(D)J recombination process in these cells. In addition, we did not observe any significant changes in lymphocytes or other peripheral blood components in the Ku70-ΔSAP mice at 180 days of age (Table S1), supporting the notion that the SAP domain is not essential for the development of lymphocytes.

### Deletion of mouse KU70-SAP domain sensitized mice to TBI

The current structural models of the DNA bound DNA-PK complex suggest that an intermediate step is involved in protecting and making DNA ends ligation-compatible before the long-range synapsis complex is converted to the short-range complex for end-ligation (4,6,24). Interestingly, two distinct long-range complexes were observed, each coping with different scenarios of end processing and ligation (6), indicating that at least two types of intermediate steps may have evolved to process different types of DNA ends. This scenario makes biological sense because the processing and re-ligation of V(D)J ends would differ from the steps required to ligate DSB ends generated by IR, which often have “dirty” and non-uniform chemical structures. Therefore, it is important to determine whether the SAP deletion affects the repair of exogenously induced DSBs, despite its insignificant impact on V(D)J recombination and lymphocyte development.

With this understanding, we sought to determine whether the deletion of the SAP domain affects KU70’s ability to manage IR-induced DNA damage at the whole-animal level.

It is well-established that mammalian hematopoietic stem cells (HSCs) and hematopoietic progenitor cells (HPCs) are among the cell types most sensitive to IR-induced DNA damage.

TBI-induced killing of HSCs and HPCs impairs hematopoiesis, which is essential for replenishing the functional pool of blood components. This impairment leads to BM failure and often results in animal death, typically occurring within 1-3 weeks post-TBI. To test whether the SAP deletion confers a sensitization of the BM to radiation damage, we exposed sex-matched littermates to an 8 Gy TBI, a dosage known to cause mouse lethality due to BM failure (25). We observed that the KU70-ΔSAP mice were hypersensitive to IR. As shown in Figure 2, about 70% of the WT mice survived the TBI, but only 20% of the KU70-ΔSAP mice did so.

**Figure 2.**
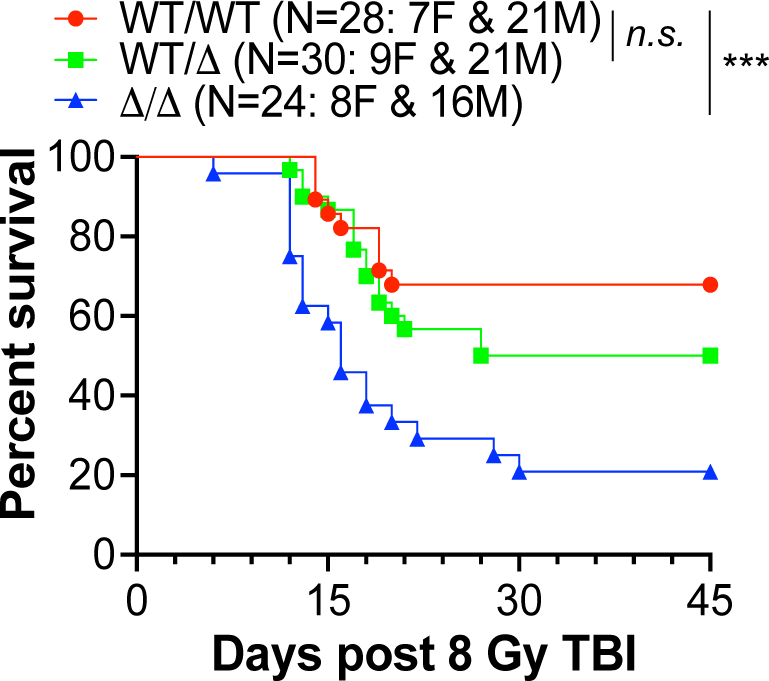
**Kaplan-Meier survival curves of KU70-**Δ**SAP mice after 8 Gy of total body irradiation (TBI).** Eight-weeks old littermates of WT (WT/WT), heterozygous KU70-ΔSAP(WT/Δ), and homozygous KU70-ΔSAP (Δ/Δ) were exposed to a single dose of 8 Gy γ- radiation by TBI. The number of male (M) and female (F) mice in each group are shown. Log- rank test was used for statistical analysis. \*\*\**p* ≤ 0.001. *n.s.* denotes non-significant.

To confirm whether the increased sensitivity of KU70-ΔSAP mice was attributed to the sensitivity of the hematopoietic stem cells to IR, we performed a hematopoietic lineage analysis using a previously established approach (17). A sublethal radiation dose of 6.5 Gy was administered to ensure the animals remained viable for BM collection and subsequent analysis, a standard approach in the field. Briefly, BM mononuclear cells were obtained from the femur, tibia and spine bones of each mouse. Among the lineage-negative (Lin^-^) cells, Sca-1 c-Kit (LS-K) and Sca-1 c-Kit (LSK) cells were further categorized based on cell surface markers using the same approach as we reported before (17). Representative flow cytometry profiles are shown for the KU70 KI mouse (Figure 3A). In the absence of irradiation, there was no difference in the total number of long-term hematopoietic stem cells (LT-HSCs), short-term hematopoietic stem cells (ST-HSCs), multipotent progenitors (MPPs), lymphoid-primed multipotent progenitors (LMPPs), common myeloid progenitors (CMPs), megakaryocyte-erythroid progenitors (MEPs), granulocyte-monocyte progenitors (GMPs) and common lymphoid progenitors (CLPs) between KU70 WT and KU70-ΔSAP mice (data not shown). However, at 14 days post IR, KU70-ΔSAP mice exhibited a notable decrease in LT-HSCs and ST-HSCs (Figure 3B). In addition, there were significant reductions in the committed progenitor populations such as CMP, MEP, and GMP (Figure 3B). The numbers of MPPs, LMPPs and CLPs did not differ significantly between KU70-WT and KU70-ΔSAP mice. These findings suggest that the SAP domain of KU70 plays a crucial role in the survival of hematopoietic stem and certain progenitor populations following IR-induced damage, as well as in the subsequent hematopoiesis required to refurnish the BM and blood.

**Figure 3.**
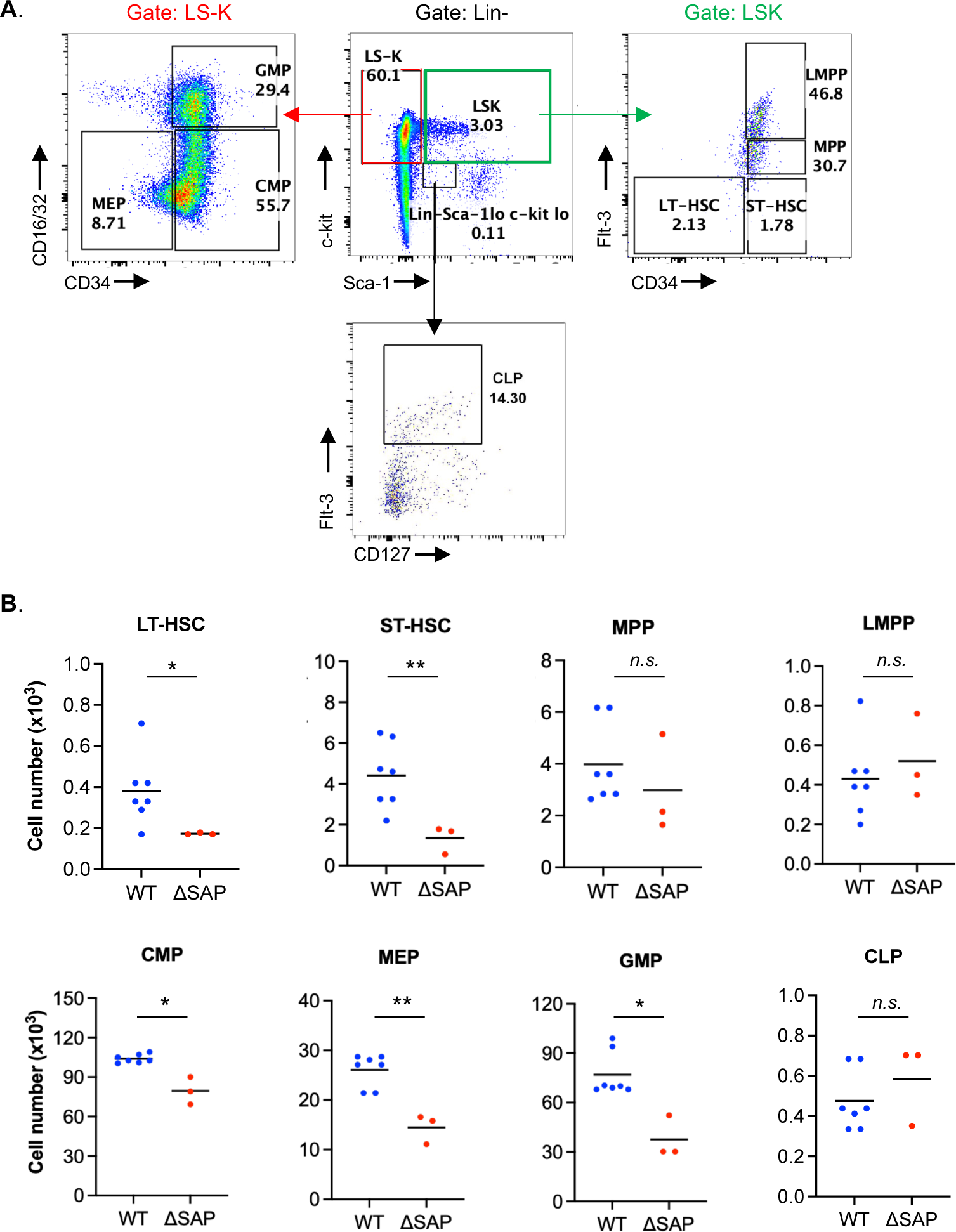
Counts of hematopoietic stem cells (HSCs) and progenitor cells in the bone marrow (BM) post TBI. Eight-week-old male mice were exposed to a sub-lethal dose of 6.5 Gy γ-radiation by TBI. The BM cells were collected at 14 days post TBI, and Lineage negative (Lin^-^) cells were isolated. (**A)** The flow cytometry gating strategy identified various cell populations: LS-K (Lin , Sca-1 , c-kit , red box) was further divided to CMPs (CD16/32 CD34 ), GMPs (CD16/32 CD34 ), and MEPs (CD16/32 CD34 ). LSK (Lin , Sca-1 , c-kit , green box) was divided to LT-HSCs (CD34 Flt-3 ), ST-HSCs (CD34 Flt-3 ), MPPs (CD34 Flt-3^Int^), and LMPPs (CD34 Flt-3^Bright^); Sca-1^low^, c-kit^low^ (black box) was used to gate the CLP population. (**B**) The total number of each cell population is presented as mean (n = 3-5 mice/group). \**p* < 0.05; \*\**p* < 0.01; and *n.s.* denotes non-significant by unpaired two-tailed Student’s t-test.

### SAP domain deletion increased IR-induced chromosome damage

Aberrations of metaphase chromosomes is a reliable indicator for defective repair of radiation-induced DNA damage. To address whether the deletion of SAP domain impairs the repair of radiation-induced DSBs, we collected splenic B-lymphocytes, stimulated their proliferation in culture, and exposed them to γ-rays. Then, metaphase spreads were prepared and scored for chromosome aberrations. As shown in Figure 4, in the absence of radiation exposure, KU70-ΔSAP B-cells exhibited approximately a 3-fold increase in spontaneous chromosome aberrations compared to the WT cells. Following exposure to 2 Gy of γ-rays, the KU70-ΔSAP cells showed a higher frequency of chromosome aberrations (Figure 4A). This increase was mostly contributed by chromosome and chromatid breaks, which are known to result from defective DSB repair, including deficiencies in NHEJ throughout the cell cycle (Table S2). Furthermore, we consistently recovered fewer metaphase cells from irradiated KU70-ΔSAP cells compared to irradiated WT cells, indicating that KU70-ΔSAP cells are more sensitive than KU70 WT cells.

**Figure 4.**
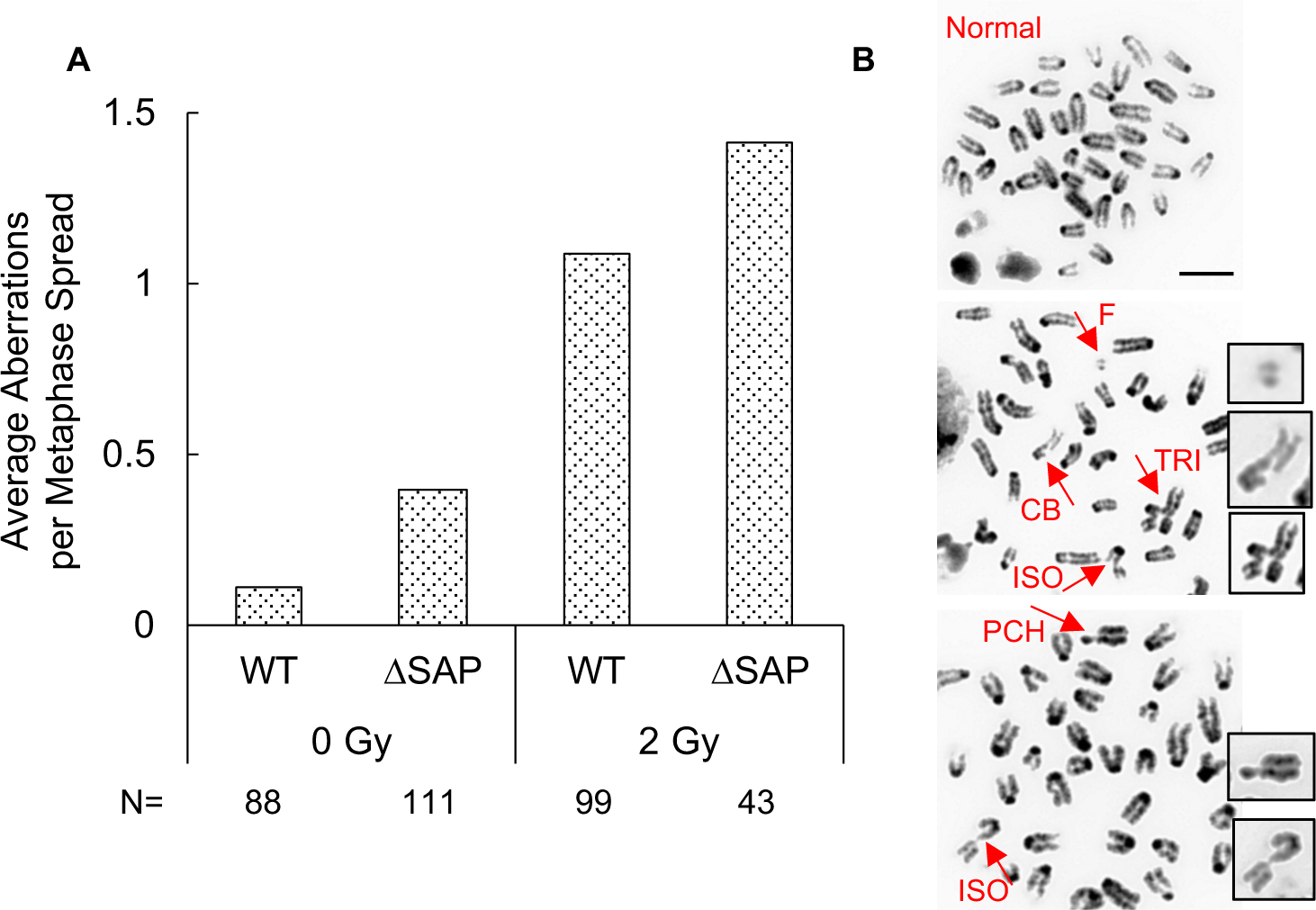
Metaphase Chromosomal aberrations in splenic B cells of KU70-WT and. Δ**SAP mice.** The activated B cells were treated with or without 2 Gy IR and then collected for metaphase spread. (**A**) The average number of chromosome aberrations in each metaphase spread. The numbers of scored metaphase spreads are indicated. (**B**) Representative image of DAPI Stained metaphase spreads with arrows indicating chromosomal aberrations. Scale bar: 7.7 μm. Abbreviations: CB: Chromatid Breaks; F: Fragment; TRI: Triradial; ISO: Isochromosome; PCH: Premature Chromatid Separation. See Table S2 for all types of aberrations scored.

### SAP domain deletion sensitized MEF to DSB-inducing chemotherapy agents

Having found that mouse KU70-SAP deletion sensitized BM stem cell to IR *in vivo* (Figures 2 and 3) and increased the frequencies of chromosome breaks in cultured B-cells (Figure 4), we developed a third assay to confirm the role of KU70-SAP in DSB repair. Immortalized Mouse Embryonic Fibroblast (iMEF) lines were established from both WT and KU70-ΔSAP mice. These iMEFs were exposed to radiation-mimicking chemotherapy agents bleomycin (Figure 5A) and phleomycin (Figure 5B), along with topoisomerase-2 poison VP16 (Figure 5C) known to induce DSBs. As shown in Figure 5, the KU70-ΔSAP cells displayed significantly higher sensitivity to these DSB-inducing agents compared to KU70 WT cells. Collectively, the results from Figures 2-5 suggest that deletion of the SAP domain leads to heightened sensitivity to exogenously induced DSBs in mice, despite its minimal impact on physiological V(D)J recombination (Figure S2C) and lymphocyte development (Table S1). These findings are promising as they suggest that the SAP domain is indeed crucial for repairing exogenously induced DSBs, while not being essential for repairing the physiologically induced DSB during V(D)J recombination (Figure S2C). This highlights the SAP domain and its related functions as potential targets to sensitize cells to therapeutic DSBs while minimizing adverse effects on the physiological processes of animal growth and immunological development.

**Figure 5.**
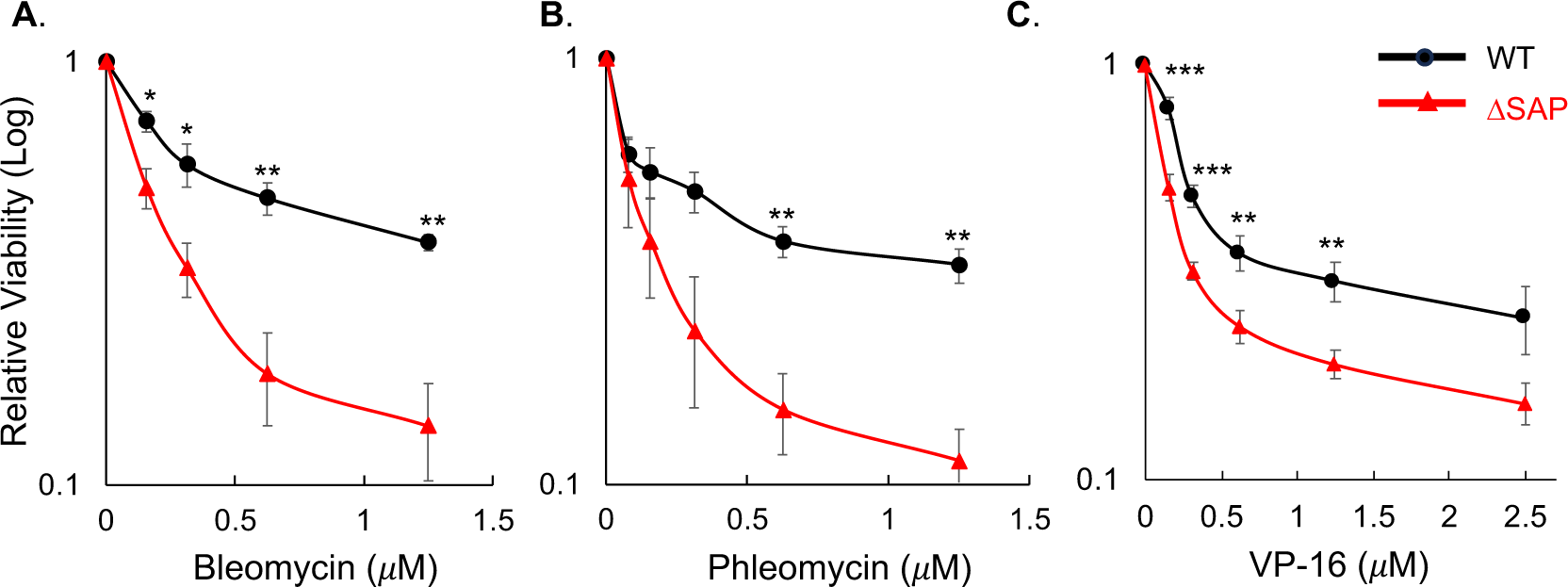
Drug sensitivity of MEFs. The MEFs were established from embryos of littermates upon cross between heterozygous KU70-**Δ**SAP mice. **The viability of each group** was recorded by the IncuCyte live-cell imager every 2 h. Shown are the relative cell viabilities at 72 hours after continuous exposure to (**A)** Bleomycin, (**B**) Phleomycin and (**C**) VP16 at the indicated concentrations, as normalized to the same cells without drug treatment. Results were the average of at least six independent experiments. Error bars are the standard errors of the mean (SEMs). P values were calculated based on two-tailed student t-test. \**p* ≤ 0.05, \*\**p* ≤ 0.01, \*\*\**p* ≤ 0.001.

### Delayed recruitment of KU70-**Δ**SAP at DNA damage sites in MEFs

To explore the cellular mechanisms by which SAP deletion confers a sensitization to exogenous DSBs, we compared the recruitments of KU70 WT and ΔSAP to DNA damage stripes after laser micro-irradiation. By eliminating RNA-associated KU70, we visualized KU70 recruitment at the irradiated nuclear stripes with minimum background signal in iMEFs (Figure 6A). Intensity quantifications of KU70 and γH2AX staining at the irradiated stripes revealed a significant reduction in KU70-ΔSAP recruitment compared to KU70 WT (Figure 6A, 6B). Furthermore, a lower percentage of cells with positive recruitment of KU70-ΔSAP than WT KU70 as compared with the γH2AX stripe-positive cells was observed (Figure 6C). For example, 15 min after micro-irradiation, KU70-ΔSAP showed only 50% of the intensity relative to KU70 WT at the irradiated stripes (Figure 6B). Additionally, only 50% of the γH2AX positive cells also exhibited KU70-ΔSAP positivity, compared to 80% of them positive for KU70 WT. Interestingly, in contrast to the reduced KU70-ΔSAP recruitment and sharpen localization at the irradiated stripes, we observed a spreading pattern of the γH2AX staining in the KU70-ΔSAP cells (Figure 6D, S3), indicating an expanded DNA damage signaling and/or defective repair, or an altered chromatin dynamics that warrant further investigation. Collectively, these findings highlight the critical role of the KU70-SAP domain in efficiently recruiting KU70 to DNA damage sites and in the repair of DSBs in the live cells.

**Figure 6.**
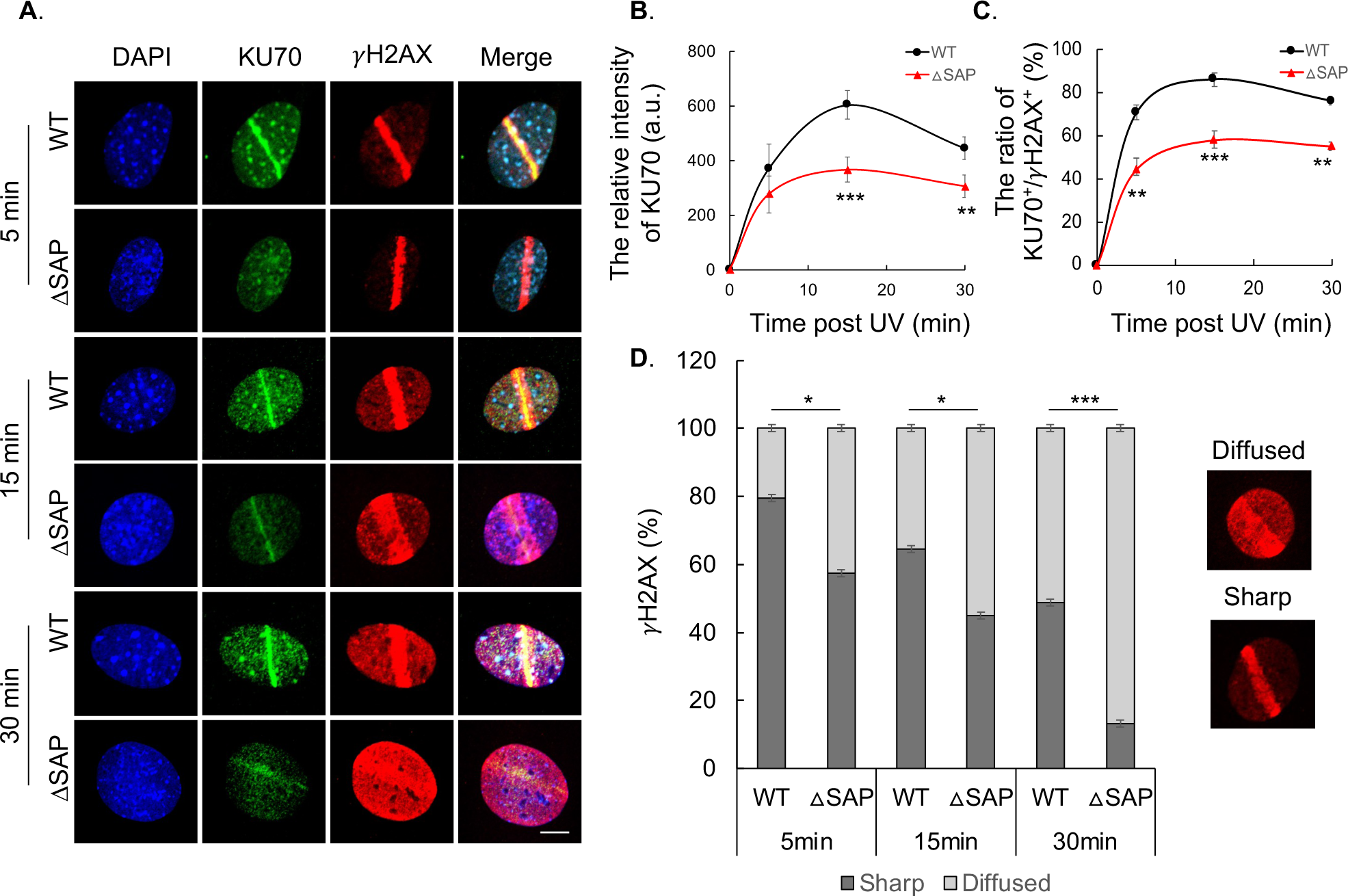
KU70 recruitment to DNA damage sites. (A) Representative confocal IF staining images of KU70 (green) and YH2AX (red) post micro-irradiation at the indicated time points. Cell nuclei were visualized with DAPI (blue). Scale bars, 10 μm. (**B**) The relative fluorescence intensity of KU70 in WT and ΔSAP MEFs at the irradiated sites were quantified at the indicated time points post micro-irradiation. (**C**) The ratio of KU70^+^ and YH2AX^+^ cells. For each time point, >40 nuclei were analyzed in three independent experiments. (**D**) The percentage of different patterns (sharp or diffused, shown on the right panel) of YH2AX in WT and ΔSAP MEFs post micro-irradiation at the indicated time points. Error bars: SEM. P values were calculated using a two-tailed Student’s t-test in (B), Binomial z statistic pairwise comparison was used for statistical analysis in (C) and (D). \**p* ≤ 0.05, \*\**p* ≤ 0.01, \*\*\**p* ≤ 0.001.

### Deletion of the KU70-SAP domain reduced the retention of LIG4 at the DSB sites

Numerous findings have provided detailed insights into crucial stages of the NHEJ pathway. According to a prevailing model, the DNA-PK complex initially binds to the broken DNA ends, attracting complex of LIG4/XRCC4/XLF. This assembly forms a synaptic complex responsible for tethering and rejoining DNA ends (2,26–29), which does not need the enzymatic activity of the LIG4 (4,6,24). Given that the SAP deletion did not affect the normal V(D)J repair process but specifically impacted exogenously induced DSB repair, our speculation is that the deletion likely interferes with a step occurring after the formation of the initial DNA-PK complex. This step is considered a fundamental process in a subset of NHEJ events. We also investigated LIG4 as a representative of the LIG4/XRCC4/XLF oligomers. Surprisingly, we noticed a diminished presence of LIG4 at the stripes during later timepoints. As shown in Figure 7, while the recruitment of LIG4 was similar between KU70 WT and ΔSAP at 5 minutes post-micro- irradiation, there was a more pronounced reduction in LIG4 intensity in KU70-ΔSAP cells compared to KU70 WT by the 15-minute mark (Figure 7). This finding implies that the SAP domain may play a crucial role in retaining LIG4 at the damage sites, even though it might not be indispensable for the initial recruitment of LIG4.

**Figure 7.**
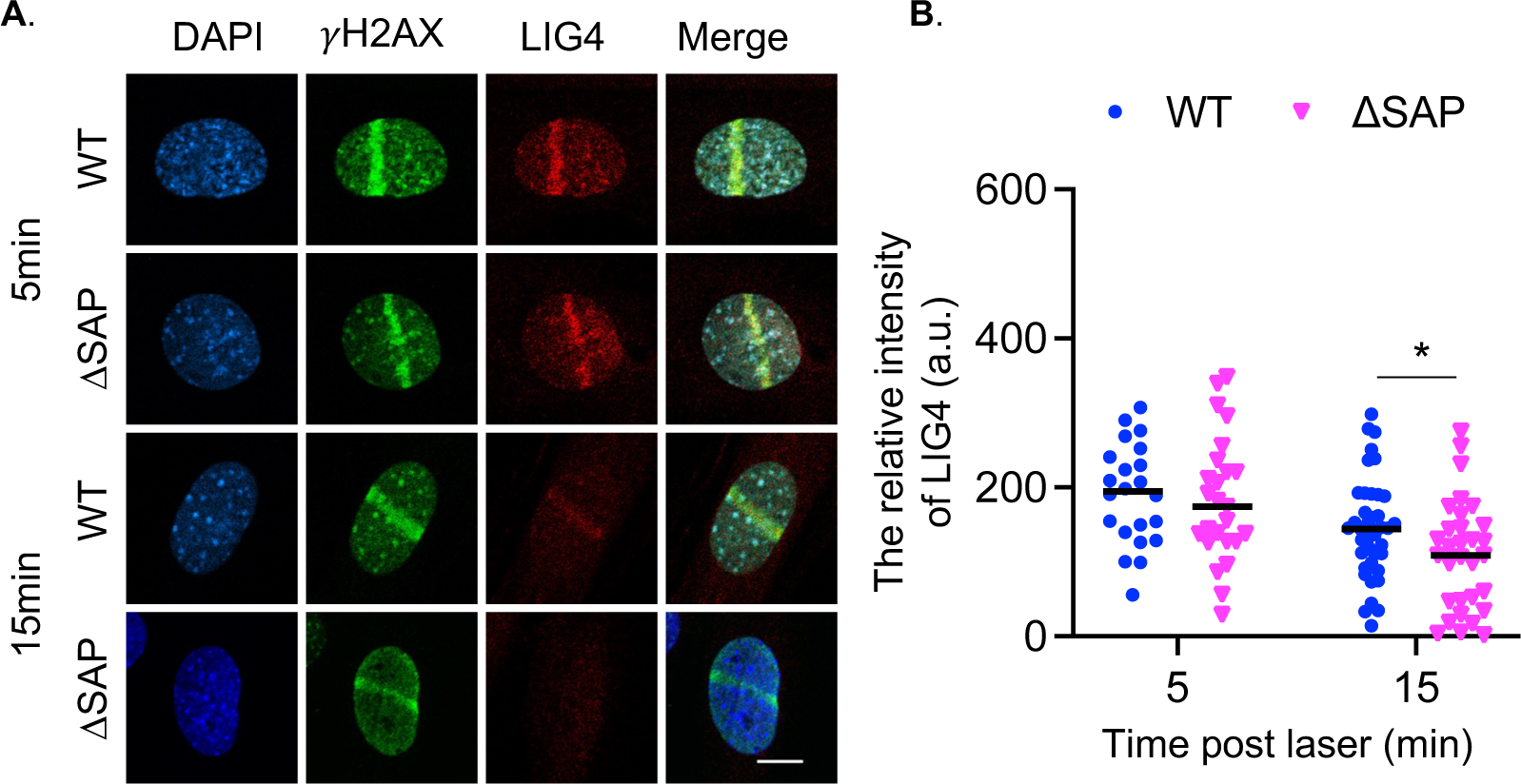
The recruitment of LIG4 to DNA damage sites. **(A)** Representative confocal IF staining images of LIG4 (red) and YH2AX (green) in KU70 WT or ΔSAP MEFs post micro- irradiation at the indicated time points. Scale bars, 10 μm. (**B**) The relative fluorescence intensity of LIG4 at the irradiated sites were quantified at the indicated time points post micro-irradiation. For each time point, >20 nuclei were analyzed in three independent experiments. Error bars: SEM. P values were calculated using a two-tailed Student’s t-test. \**p* ≤ 0.05.

### Lack of SAP domain dampened LIG4 and other NHEJ cofactors chromatin-binding in human cells

To confirm the reduced retention of LIG4 at DNA damage sites, a human KU70 knockdown system was implemented. Recognizing the essential role of human KU70 in cell viability, an inducible KU70 knockdown strategy was employed to investigate the impact of KU70-SAP in human cells. Utilizing the H1299 cell line, we established a doxycycline (Dox)-inducible KU70 knockdown system. Subsequently, we reintroduced various exogenous human KU70 variants, including KU70-WT, KU70-ΔSAP, and KU70-SAP (comprising both the nSAP and cSAP domains of KU70) (Figure 8A). The expressions of these variants and their nuclear localization were verified in the cells using Myc-EGFP tag (Figure S4).

**Figure 8.**
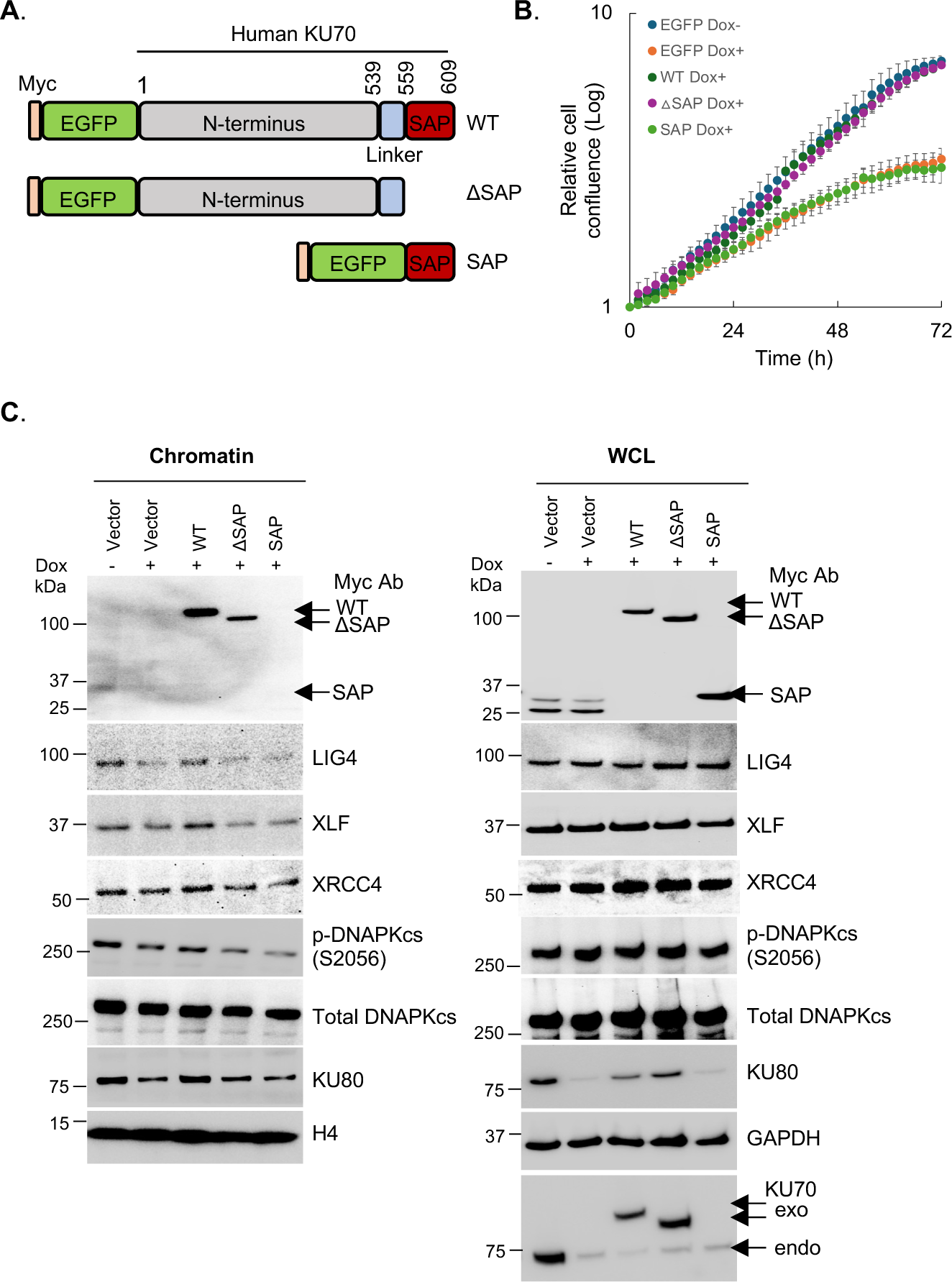
**KU70-**Δ**SAP reduced recruitment of NHEJ co-factors to chromatin in the H1299 cell line following IR.** (**A**) Schematic representation of Myc-EGFP tagged full length KU70 (WT) and two truncated mutants (ΔSAP and SAP). (**B**) Growth curve of KU70 knockdown cells stably expressing Myc-EGFP-tagged KU70 (WT and mutants). Following 3 days treatment with Dox to induce the knockdown of endogenous KU70, cells were re-plated to monitor cell growth for a period of 72 hours. The Dox- served as the control to compare with endogenous KU70 knock down. Error bars: SEM. (**C**) Western blot analysis of chromatin bound fractions from H1299-shKU70 cells treated with or without Dox, stably expressing Myc-EGFP-tagged KU70 (WT and mutants), 30 min post 6.5 Gy IR. WCL: whole cell lysate. The blots were probed with indicated antibodies.

As anticipated, the exogenous expression of KU70 WT fully restored the proliferation defect induced by endogenous KU70 depletion following Dox treatment. Interestingly, KU70-ΔSAP showed a growth rate similar to KU70-WT, indicating that the deletion of the human SAP domain did not impact cell growth (Figure 8B), aligning with the normal development observed in KU70-ΔSAP mice (Figure S2). However, the expression of the SAP domain alone failed to rescue cell viability, underscoring that the SAP domain itself is insufficient to sustain cell viability despite being localized in the nucleus (Figure S4).

We then assessed the chromatin-bound fraction of these KU70 protein variants. As shown in Figure 8C, Dox-induced knockdown of endogenous KU70 significantly decreased the levels of chromatin-bound LIG4, XLF, XRCC4, KU80, and p-DNA-PKcs at S2056 compared to cells without knockdown (Dox-). However, the chromatin binding was largely restored upon re-expression of Myc-EGFP-tagged WT KU70. Reintroduction of KU70-ΔSAP and KU70-SAP did not restore the levels of these chromatin-bound NHEJ factors to the same level as KU70 WT, despite the total protein levels being similar to KU70 WT cells (Figure 8C). It is interesting that KU70-ΔSAP cells exhibited a notable reduction in chromatin-bound LIG4 compared to cells expressing KU70 WT, as well as LIG4 retention at the micro-irradiated DNA damage sites in the MEF cells (Figure 7), emphasizing the role of the KU70-SAP domain in preserving LIG4 on chromatin post DNA damage. While there were reductions of chromatin bound KU80 and p- DNA-PKcs (S2056) after irradiation of the human cells, this was not obvious in micro-irradiated nuclear stripes in the MEFs.

Together, our data (Figure 7 and 8) strongly suggest that the deletion of the SAP domain results in a less sustainable association of LIG4 with the DNA repair complex in the cells, offering a novel mechanism for KU70, especially its SAP domain, to regulate a critical step of NHEJ after formation of the DNA-PK complex.

## Discussion

Despite previous efforts to elucidate the structural characteristics of KU70-SAP and the biochemical understanding that the KU70-SAP may bind to RNA or DNA (14–16), the specific cellular function of the mammalian KU70-SAP has not been documented in the literature. In a recent study (15), the *Arabidopsis thaliana* KU70-SAP was found to facilitate the DNA end loading of KU dimer but it was not required for classical NHEJ and telomere maintenance in the plants. Because the SAP deletion in the *Arabidopsis thaliana* study retained the nSAP domain (15), in addition to that *Arabidopsis thaliana* does not have a DNA-PKcs (5) and thus may use a distinct KU-dependent mechanism to repair DSBs, it is of particular interest for DNA repair and cancer research fields to directly define the function of mammalian KU70-SAP.

In this study, we investigated the deletion of both the KU70-specific nSAP and the generically defined cSAP of mouse KU70 and found that these deletions did not affect the gross animal development and the physiological V(D)J recombination process. However, they did have a severe effect on animal sensitivity to IR, chromosome aberrations, and cellular sensitivity to exogenously induced DSBs. We further showed that SAP deletion led to a decreased efficiency of the KU70-ΔSAP in reaching DNA damage sites in cells, Additionally, there was an early departure of LIG4 from the DNA damage sites associated with SAP deletion. Our discoveries are significant as they indicate that the SAP domain plays a vital role in cellular repair of exogenous DSBs, while being less critical for physiological processes. Therefore, targeting the SAP domain for inhibition could potentially enhance cell sensitivity to radiation and other therapeutic agents that induce DSBs.

Recently, there have been major breakthroughs in the structural understanding on how the DNA-PK complex and associated dynamic proteins execute the ligation of two dsDNA ends (6,10). The consensus model is that following the encircling of dsDNA by the KU dimer and the recruitment of DNA-PKcs to protect the DNA end, the KU dimer shifts inward from the break end to stabilize the DNA (30–32). However, there are alternative configurations of protein complex after the initial DNA-PK complex is established. For examples, this complex can recruit LIG4/XRCC4/XLF and other proteins to form either domain-swap or XLF-mediated conformations, each with distinct activities tailored towards different types of DNA ends (6).

Therefore, it is plausible that distinct evolutionary steps have been developed to address various DNA strand breaks, and a defect in one specialized conformation may not necessarily impact the function of another, potentially resulting in different repair outcomes based on the types of DSBs. Our results suggest that SAP domain is essential for a specific conformation that favors the repair of exogenously induced DSB, but it is not required for the ones to perform V(D)J recombination. This highlights the intricate and specialized roles that different configurations of the protein complex play in DNA repair processes, depending on the nature of the DNA damage encountered.

The next question of why and how KU70-SAP is required for repairing of exogenously induced DSBs but not for V(D)J recombination prompts a deeper exploration. One distinction between DSBs generated during the V(D)J and those induced by exogenous insults is the chemical nature of the break ends. The V(D)J ends are processed by cleavage of the hairpin-loop with enzymes that can directly result in the ligation compatible ends. In contrast, the IR and chemotherapy induced DSBs are likely to have un-uniformed chemical structure at the ends, which would require extra-end processing steps before they become ligation compatible. This may require an extra regulatory step that is fulfilled by the SAP domain. However, the specific mechanism by which SAP deletion renders cells sensitive to exogenous DSB, but not impact V(D)J related to strand breaks, would need further investigation.

Delving into a more detailed analysis, the present study has provided new support for a conceptual refinement of the KU70-SAP domain that holds functional significance. As illustrated in Figure S1, the commonly referred KU70-SAP is composed of two distinct sub-domains. It is only the cSAP that aligns with the originally defined SAP motif of approximately 35 aa, but the 14 aa nSAP forms a new helix that appears to have evolved specifically for KU70, although the nSAP and cSAP together form a globular structure together. Interestingly, among all possible positions of the KU70-SAP, it is the nSAP helical domain that serves as the interaction interface between the KU70-SAP and the main body of the KU dimer (7,8,13). Thus, we speculate that the KU70-nSAP is likely evolved to mediate certain KU70 specific function via its interaction with the KU dimer and others, while the cSAP fulfills the more generic functions commonly associated with SAP domains. This concept emphasizes the complex interaction dynamics between the distinct sub-domains of the KU70-SAP, underscoring their specific functions in facilitating precise protein interactions within the DNA repair complex.

The current results further suggest that the SAP domain achieves these functions via regulating LIG4 and also likely its associated partners which could impact its ligase activity. This regulatory function is particularly relevant during the transition of LIG4 from a non-enzymatic role in tethering DNA ends within the long-range synapsis complex to a state ready for ligation within the short-range synapsis complex. This transition likely necessitates a reconfiguration of LIG4 to enable its flexible ligase domain to access the ends that are suitable for ligation. Coincidentally, the nSAP helix were shown to dock around the α/β-domain of KU80 (7,8) or the KU aperture area (13), which overlap with the KU80 core region and the KU70 C-helical ARM that bind to the LIG4 (4), while the cSAP has not been implicated in any of these interactions.

LIG4 retention was consistently reduced at micro-irradiated stripes as well as in the chromatin-bound form in irradiated cells (Figure 7, 8C). Our observation may indicate a role of the SAP domain in the stability of the short-range complex through the retention of LIG4, a prediction requiring additional experimental testing. LIG4 has at least two roles in the NHEJ. First, two LIG4, two XRCC4, and a XLF homodimer form the long-range synapsis complex to tether two DNA ends. This function does not need the ligase activity (24) but is still critical for NHEJ. When transitioning to the short-range complex, the flexible ligase domain of LIG4 must be reconfigured to access the synapsed DNA ends, allowing each LIG4 molecule to ligate one strand and facilitate the rejoining of the DSB ends. Interestingly, this LIG4 enzymatic activity appears be dispensable for NHEJ, likely due to LIG3 or other yet-to-be-identified compensating ligase activity (33). Because SAP loss impaired LIG4 retention, which would also call for an alternative ligase to rejoin the break as proposed by Golf et al (33), our results hint at a potential synthetical lethal relationship between the SAP domain and the putative LIG4 backup ligase during the NHEJ. Furthermore, it is noteworthy that the nSAP binding interfaces on the KU dimer appear to overlap with some of the binding surface of LIG4. Thus, it is possible that the SAP domain may coordinate with LIG4 to reconfigure the flexible ligase domain to promote end-ligation. This prediction is consistent with the structural model where the non-enzymatic LIG4 is crucial to form and maintain the long-range and short-range synapsis complex, but the enzymatic domain of LIG4 is reconfigured to localize and ligate one strand of the DNA, which seems compensable by other ligase(s) when the enzymatic activity of LIG4 is inactivated.

Overall, the present findings are exciting as they have demonstrated that SAP deletion conferrs a sensitivity to exogenous DNA damage from irradiation and DSB-inducing chemotherapy agents, while having no apparent impact on physiologically induced DSBs, such as those involved in V(D)J recombination. This suggests that specifically disrupting SAP-related function could be a promising strategy to enhance cell sensitivity to therapeutic DNA damage, while minimizing adverse effects on the physiological activity of KU70 in animal development and immunological response. Our functional and cell biology analyses also raise interesting questions for biochemical and structural refinement of the NHEJ process, as well as open new opportunities to manipulate the NHEJ process for therapeutic development.

## Supplementary data

Supplementary Data are available Online

## Acknowledgements

We thank Dr. Yantao Zuo (Rutgers Cancer Institute) for critical reading of the manuscript before submission, Dr. Sam Bunting (Department of Molecular Biology and Biochemistry, School of Arts and Sciences, Rutgers University) for providing technical consultation on class-switch analysis.

## Author contributions

YW designed the experiments, performed data analysis, and drafted the manuscript. YW, MC, HK, HL, JL, YC, and PR performed experiments. CM provided resources for the experiments and advise on experiment protocols and scored chromosome aberrations. ZS co-designed the experiments, directed and oversaw the project, secured funding for the study, and completed the final version of the manuscript.

## Declaration of data availability

The data underlying this study will be deposited to public database per NIH policy and can be made available to readers.

## Supplememtal Figures

**Figure S1. Distinction between the canonical SAP (cSAP) and the nSAP domains of KU70.** While the cSAP domains of human and mouse KU70 align with the originally defined SAP domain, the nSAP domain appears to be specific to KU70.

Figure S2. Absence of significant spontaneous developmental defects and tumor burden in Δ**SAP mice.** (**A**) The body weight of WT and ΔSAP mice was recorded over the first 3 months after birth. (**B**) The distribution of mice with or without the tumors was assessed in both WT and ΔSAP mice. Mice were sacrificed at 600 days after birth to evaluate the presence of spontaneous tumors or other abnormalities. (**C**) Immunoglobulin class switching in stimulated B cells was measured by the frequency of IgG1. The left two panels show representative flow cytometric profiles of IgG1 conversion from IgM in WT and ΔSAP mice. The right panel summarizes the frequency of IgG1 in IgM^+^ cells based on three independent experiments. P values were calculated using a two-tailed Student’s t-test. *n.s.* denotes non-significant.

**Figure S3. Representative sharp or diffused pattens of** γ**H2AX post micro-irradiation at the various time points in KU70-WT and** Δ**SAP cells.** The intensity of γH2AX along a line crossing the stripe is shown in the chart on the left.

**Figure S4. Intra-cellular localization of Myc-EGFP tagged KU70 (WT and mutants) in H1299 cells.** Cell nuclei were visualized with DAPI (blue). Scale bars, 40 μm.

