## Supplementary figures and images for "The Mammalian KU70 C-terminus SAP Domain Is Required to Repair Exogenous DNA Damage"

### Supplement Figures 1-4

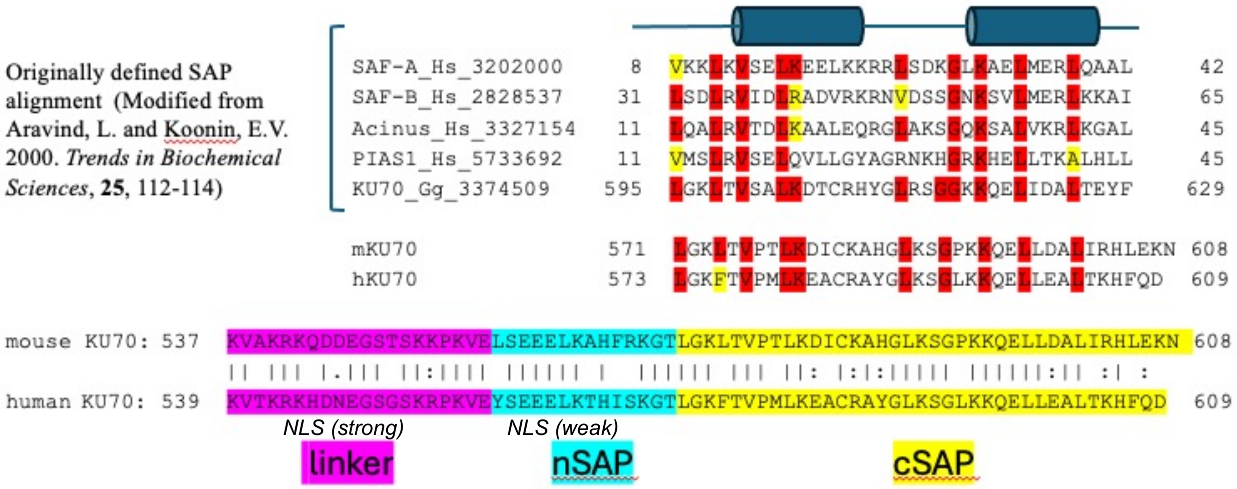

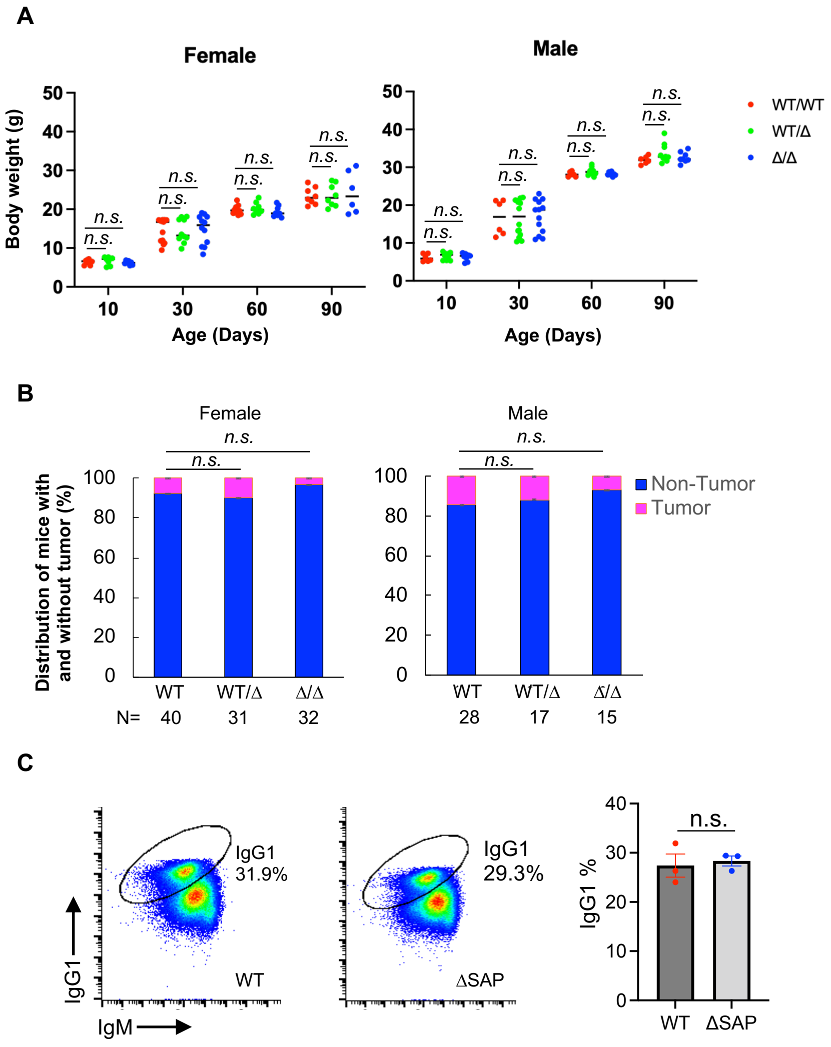

#### Figure S3

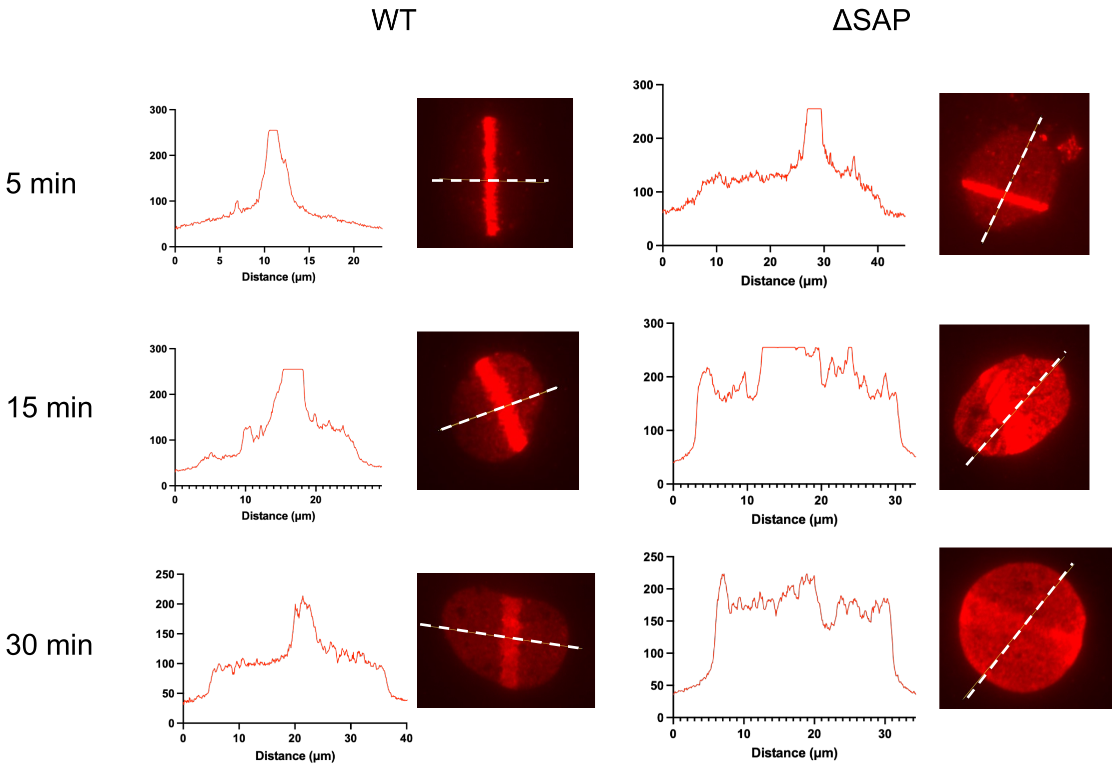

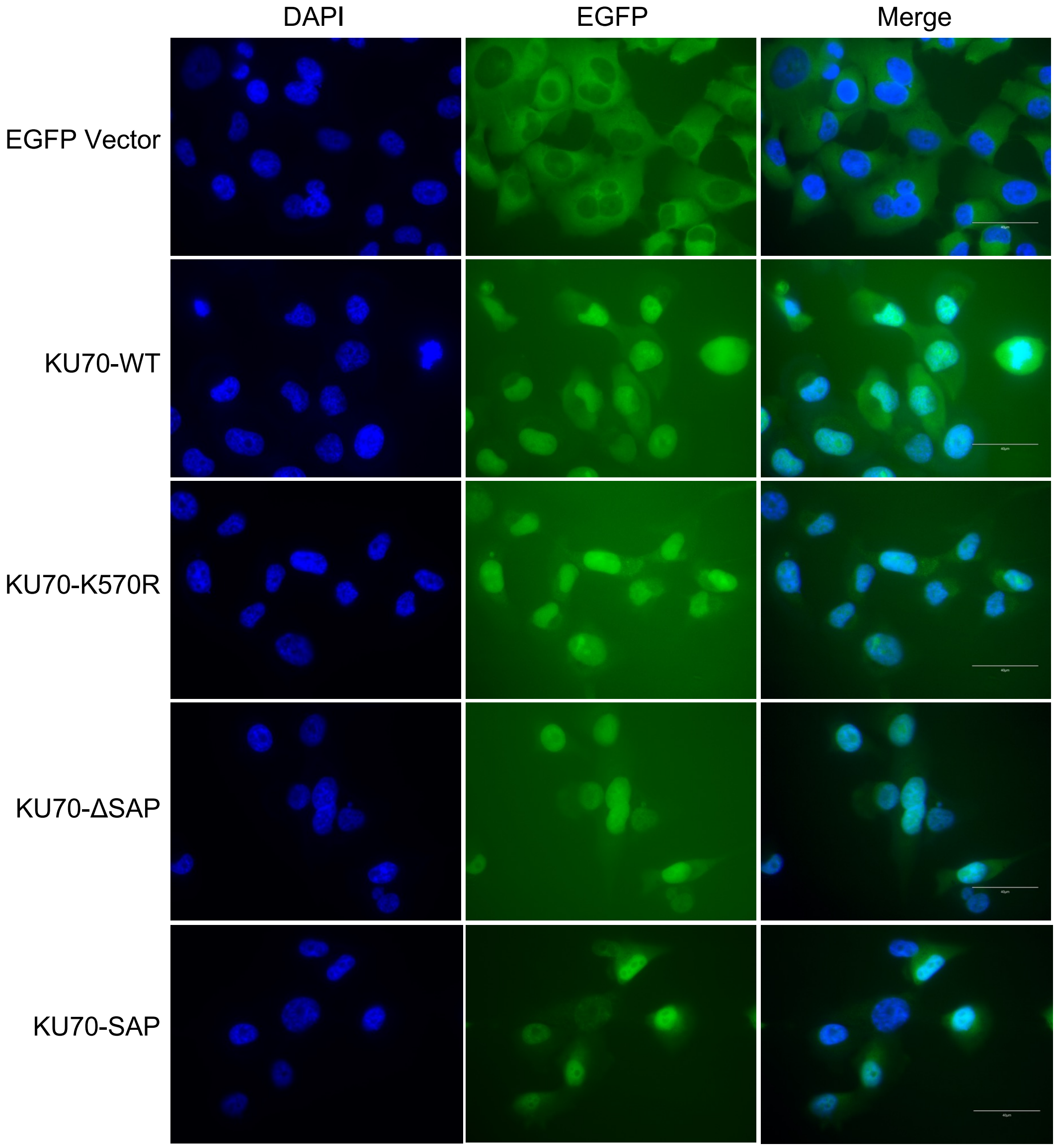
