## Supplement tables S1 and S2 for "The Mammalian KU70 C-terminus SAP Domain Is Required to Repair Exogenous DNA Damage"

**Table S1. Complete blood count from KU70-WT and ΔSAP Mice.** Blood samples were drawn from the submandibular vein of 180-day old mice (N=3-5), collected in BD vacutainer EDTA tubes (#367844) and analyzed with the HESKA-HT5 veterinary hematology analyzer. No significant statistical differences were observed in the results. Abbreviations: WBC, white blood cells; Neu, neutrophils; Lym, lymphocytes; Mon, monocytes; Eos, eosinophils; Bas, basophills; RBC, red blood cells; HGB, hemoglobin; HCT, hematocrit; MCV, mean corpuscular volume; PLT, platelets; MPV, mean platelet volume.

| Parameter | WT | ΔSAP |
|---|---|---|
| WBC ($10^3/\mu L$) | 10.20 ± 0.21 | 9.34 ± 1.32 |
| Neu (%) | 11.45 ± 0.49 | 13.98 ± 1.40 |
| Lym (%) | 86.38 ± 0.05 | 82.73 ± 1.33 |
| Mon (%) | 1.28 ± 0.41 | 1.57 ± 0.12 |
| Eos (%) | 0.51 ± 0.11 | 1.03 ± 0.18 |
| Bas (%) | 0.37 ± 0.03 | 0.68 ± 0.02 |
| RBC ($10^6/\mu L$) | 9.35 ± 0.56 | 10.14 ± 0.07 |
| HGB (g/dL) | 14.26 ± 0.91 | 14.90 ± 0.06 |
| HCT (%) | 42.16 ± 2.43 | 45.13 ± 0.22 |
| MCV (fL) | 45.10 ± 0.38 | 44.45 ± 0.09 |
| PLT ($10^3/\mu L$) | 1207.67 ± 153.59 | 1419.67 ± 170.57 |
| MPV (fL) | 5.47 ± 0.06 | 5.43 ± 0.03 |

**Table S2. Summary of metaphase chromosomal abnormalities in KU70-WT and ΔSAP mouse splenic B cells, with or without IR treatment.**

| Aberration Types | 0 Gy | | 2 Gy | |
|---|---|---|---|---|
| | WT | ΔSAP | WT | ΔSAP |
| Premature Centromeric Separation (PCS) | 1 | 4 | 15 | 12 |
| Dicentric Chromosomes (DC) | 1 | 0 | 9 | 4 |
| Chromatid Break (CB) | 4 | 20 | 32 | 19 |
| Fragment (F) | 2 | 5 | 4 | 5 |
| Acentric Chromosome (AC) | 1 | 0 | 1 | 3 |
| Translocation (T) | 0 | 3 | 0 | 0 |
| Sister Chromatid Fusion (SCF) | 0 | 6 | 4 | 0 |
| End To End Fusion (ETE) | 0 | 0 | 1 | 1 |
| Quadriradial (QUA) | 0 | 0 | 0 | 1 |
| Triradial (TRI) | 1 | 0 | 3 | 3 |
| Premature Chromatid Separation (PCH) | 0 | 2 | 4 | 3 |
| Isochromosome (ISO) | 0 | 4 | 35 | 10 |
| Total Number of Aberrations | 10 | 44 | 108 | 61 |
| Total Spreads | 88 | 111 | 99 | 43 |
| Average Aberrations per Metaphase Spread | 0.11 | 0.40 | 1.09 | 1.42 |
